# Combining model-based and data-driven models: an application to synthetic biology resource competition

**DOI:** 10.1101/2025.03.09.642275

**Authors:** Atefe Darabi, Zheming An, Muhammad Ali Al-Radhawi, William Cho, Milad Siami, Eduardo D. Sontag

**Affiliations:** Department of Electrical and Computer Engineering, Northeastern University, Boston, MA, USA; Division of Genetics and Genomics, Department of Pediatrics, Boston Children’s Hospital, Boston, MA, USA; Harvard Medical School, Boston, MA, USA; Broad Institute of MIT and Harvard, Cambridge, MA, USA; Departments of Electrical and Computer Engineering and Bioengineering, North-Eastern University, Boston, MA, USA; Department of Bioengineering, Northeastern University, Boston, MA, USA; Departments of Electrical and Computer Engineering and Bioengineering, and affiliate of Departments of Mathematics and Chemical Engineering, Northeastern University, Boston, MA, USA

## Abstract

This work explores the integration of machine learning (ML) and mechanistic models (MM). While ML has demonstrated remarkable success in data-driven modeling across engineering, biology, and other scientific fields, MM remain essential for their interpretability and capacity to extrapolate beyond observed conditions based on established principles such as chemical kinetics and physiological processes. However, MM can be labor-intensive to construct and often rely on simplifying assumptions that may not fully capture real-world complexity. It is thus desirable to combine MM and ML approaches so as to enable more robust predictions, enhanced system insights, and improved handling of sparse or noisy data. A key challenge when doing so is ensuring that ML components do not disregard mechanistic information, potentially leading to overfitting or reduced interpretability. To address that challenge, this paper introduces the idea of Partially Uncertain Model Structures (PUMS) and investigates conditions that discourage the ML components from ignoring mechanistic constraints. This work also introduces the concept of embedded Physics-Informed Neural Networks (ePINNs), which consist of two loss-sharing neural networks that seamlessly blend ML and MM components. This work arose in the study of the context problem in synthetic biology. Engineered genetic circuits may exhibit unexpected behavior in living cells due to resource sharing. To illustrate the advantages of the ePINNs approach, this paper applies the framework to a gene network model subject to resource competition, demonstrating the effectiveness of this hybrid modeling approach in capturing complex system interactions while maintaining physical consistency.

## 1. Introduction

### 1.1. Machine learning or/and mechanistic models?

Machine learning (ML) approaches have advanced rapidly in recent years, demonstrating remarkable success in extracting complex patterns from large datasets and driving innovation across numerous fields [1, 2]. In systems and synthetic biology, in particular, ML can be integrated into genetic circuit design, providing new approaches to designing and optimizing synthetic biological constructs [3]. However, purely data-driven ML models often lack explanatory power and can struggle in scenarios where data is limited or do not adequately capture the underlying physics or biology of a process [4]. In contrast, mechanistic models (MM) are grounded on established principles, such as fluid dynamics equations, chemical reaction kinetics, or physiological processes. In systems and synthetic biology, “physics-based” models play a key role when engineering cellular control systems [5]. While they offer interpretability, reliability, and the ability to extrapolate beyond observed conditions [6], such models can be labor-intensive to construct, and necessarily rely on simplifying assumptions that will not always capture the full complexity of real-world systems.

Combinations of ML and MM hold the promise of leveraging the strengths of both approaches while mitigating their respective weaknesses [7, 8]. By combining domain knowledge with data-driven algorithms, it should be possible to develop hybrid models that benefit from the interpretability and theoretical rigor of mechanistic frameworks, as well as the adaptability and predictive power of ML methods. Ideally, this synergy will enable more robust predictions, improved insight into system behavior, and the capacity to handle sparse or noisy datasets more effectively. As such, the emerging field of hybrid modeling has garnered increasing interest, with potential applications from systems biology to engineering, physics, and chemistry. To draw upon the best features of both approaches, various versions of “hybrid” ML/MM combined models have been proposed, at least since the mid-1990s, particularly in chemical process control [9, 10]. In systems biology, hybrid models have been used more recently to deal with unknown delays in transcription as well as with unknown portions of signaling pathways [11, 12, 13]. Hybrid techniques aim to synergize interpretable models based on physicochemical and biochemical knowledge with data-driven statistical learning. A recent review explores and compares approaches to build ML/MM combined models in systems biology [14].

Typically, these previous works have used ML to fit neural network models to physical constraints, or to generate data for either mechanistic or ML models to train from the other. Here, we are interested in exploring a somewhat different aspect, namely how to incorporate *partially uncertain* MM as a part of larger systems which also contain totally unknown components. We emphasize “partially uncertain” because those mechanistic parts of larger models that are perfectly known may in principle be incorporated *a priori* into a model. In this process, the subsequent ML learning phase can in effect “subtract” that known part of the model and fit the residual. (Implementing that approach is not straightforward, and combining ML with perfectly known systems is of great practical interest in itself.) Let us discuss more precisely the problem to be studied.

### 1.2. Partially uncertain model structures

For concreteness, in this work we deal with systems defined by ordinary differential equation (ODE) models of the following general form:

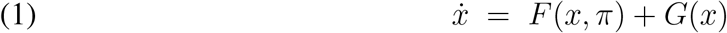

but similar considerations apply to partial differential equation, delay-differential, discrete time, or even stochastic formulations. Here *x* = *x*(*t*) is a vector function of time, taking values in ℝ ^*n*^, and dot indicates time derivative. The first term, a parameterized vector field *F*, depends on *x* as well as on a finite set of parameters *π*. It represents a “mechanistic gray box” part of the system, which is perfectly known *except* for the values of the parameters *π*. (Examples of such parameters might be binding rates in biochemistry, flow rates between body tissues in pharmacology, masses in mechanics, or resistances in electrical circuits.) This is the “interpretable” part of the system. For example, we may have a two-dimensional differential equation whose coordinates describe the position and velocity of a spring for which mass and stiffness constants are not known (the “known unknowns”). The second term, *G*(*x*), denotes a “black box” part of the system, about which nothing is *a priori* known (the “unknown unknowns”). Examples of such might be gene transcription or epigenetic effects in cell biology, immune components in mammalian organisms, or interactions among species in ecology. In the spring example, *G*(*x*) might represent unmodeled loads, nonlinear friction, or fatigue effects, thus introducing a “correction” or “compensating” term to the physically known mechanism. We will assume that *G* can be represented by a function belonging to a typically infinite dimensional “universal” function approximation class, such as deep neural networks or other machine learning architectures. One wishes to use experimental data in order to simultaneously find the (usually physically meaningful and interpretable) parameters *π* and a function *G* so that (1) is consistent with this data.

Now, one immediate problem with this admittedly vague formulation is that there is no obvious way to avoid the *overcorrection* phenomenon, in which the function *G* just chooses to *ignore all mechanistic information*. Indeed, suppose that (1) has been found to match experimental data, for a given parameter vector *π*. Now pick any other parameter 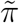. Then the model

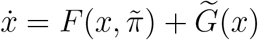

works equally well, where 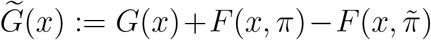. This is true even if the functions *F* (*x, π*) and 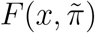 are distinct, so the issue is not one of identifiability (different parameters giving rise to the same function). In other words, using mechanistic information confers no advantage compared to a straightforward ML approach. Thus, in order for the problem to be well-posed, we need to impose some *structure* to the uncertainty, as we will discuss in Section 2.4. First, however, we will discuss a case study.

### 1.3. Context in synthetic biology

Our work was motivated by the *context problem* in synthetic biology, meaning that engineered genetic circuits may not perform as expected when inserted into living cells. Synthetic biology seeks to engineer biological systems for applications such as biosensing, programmable probiotics, and regenerative medicine. However, a significant challenge in this field is compositional context, the phenomenon where the properties of a biological circuit module change depending on the presence of other modules. This context-dependence leads to unpredictable behavior, making it difficult to design modular and scalable biological systems. Unlike traditional engineering disciplines, where standardized components interact through well-defined physical principles and abstraction layers, biological modules interact in complex and often unknown ways.

Factors such as retroactivity, resource sharing, genetic context effects, growth rate feedback, and off-target interactions can disrupt intended circuit functions [15, 16, 5] and even result in paradoxical effects in which activators of gene expression in effect function as repressors [17]. Moreover, unlike in other engineering fields where methods like feedback controllers can insulate modules from external influences, synthetic biology lacks robust frameworks to manage these context effects. As a result, genetic circuits are often designed in an ad hoc manner, requiring labor-intensive re-characterization whenever a new component is introduced. This trial-and-error approach slows down progress and limits the scalability of synthetic biology. Understanding context effects is therefore crucial for developing a more systematic and predictable design framework that accounts for unintended interactions and enables the reliable composition of biological modules. Interestingly, such context effects appear to also play a central role in naturally occurring, not synthetic, cellular processes such as for example those involved in epigenetic regulation of metastasis in cancer [18].

We will use a resource-sharing model from synthetic biology as a case study to illustrate our ideas. Specifically, we will work with the four-dimensional model presented in Section 2.2, which is a simplified version of a more detailed model also described there.

### 1.4. An approach: embedded Physics-Informed Neural Networks (ePINNs)

*Physics-Informed Neural Networks (PINNs)* are deep learning models that integrate physical laws, for example expressed as differential equations, into the training process. Unlike traditional neural networks that rely solely on labeled data, PINNs embed governing equations as constraints in the loss function. This approach has been successfully applied in fields such as fluid dynamics, material science, and biomedical engineering, where data is sparse or expensive to obtain [19]. By leveraging automatic differentiation and modern optimization techniques, PINNs provide a mesh-free alternative to conventional numerical solvers while maintaining physical consistency [20]. PINNs build upon earlier work such as [21], which trained neural networks to approximate PDE solutions by minimizing residuals. PINNs extend this idea by incorporating recent advances in deep learning to enhance accuracy, scalability, and robustness.

In this work, we extend the PINNs idea, and develop the concept of *embedded PINNs (ePINNs)*, originally proposed in [22, 23], which is a novel framework composed of two loss-sharing neural networks designed to improve predictions of unknown dynamics from quantitative timeseries data. We believe that ePINNs should be useful in the analysis of biological systems that are partially understood, addressing unknown interactions, hard-to-measure reaction rates, and missing data. Section 2.1 describes ePINNs, and we later apply ePINNs to the gene circuit resource-sharing problem.

### 1.5. Outline of paper

The paper is organized as follows. In the methods Section 2 we introduce the biological example to be used as a case study of resource competition in synthetic biology, as well as the ePINNs method and related algorithms and loss function design. We also introduce there a theoretical framework to address the overcorrection problem. The results Section 3 presents simulations, applying ePINNs to infer unknown reaction rates and predict unknown dynamics for the case study, and providing a mathematical result that helps to understand why our example is a good candidate for ePINNs modeling.

The paper ends with a discussion in Section 4.

## 2. Methods

### 2.1. Definition of embedded Physics-Informed Neural Networks (ePINNs)

In general, we will consider systems as in (1), together with a fixed choice of initial state *x*(0) = *x*_0_ and an output *y* = *h*(*x*). The measurable variables *y* = (*y*_1_, …, *y*_*l*_) are related to the system states through an output function *h* : ℝ^*n*^ → ℝ^*l*^. In typical applications, *h* is linear. We denote the *n*-dimensional Euclidean real space by ℝ^*n*^, and the set of *n* × *m* matrices with real elements by ℝ^*n*×*m*^. For a matrix *M*, its transpose is denoted as *M* ^⊤^.

As discussed in Section 1.4, this work relies upon embedded PINNs (ePINNs). We introduce ePINNs in this section.

ePINNs combine two neural networks. The first network, which we will denote by 𝒳, is parameterized by a vector of weights *θ*_1_. This network attempts to build an estimate of the state trajectory *x*(*t*) as a function of the time *t* and the initial condition *x*_0_. We write the estimated solution 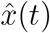 as 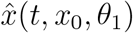 when we want to be explicit about the dependence on *x*_0_ and *θ*_1_. The second network, which we denote by 𝒢, is parameterized by *θ*_2_, and it is intended to model the unknown component *G*(*x*) of the dynamics. In summary, the system in (1) will be approximated as follows:

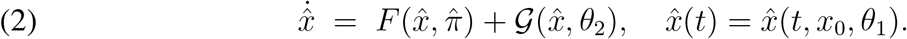

We collect the three vector parameters to be estimated into Θ = (*π, θ*_1_, *θ*_2_).

The structure of ePINNs is depicted in Figure 1. It represents a modular architecture consisting of two interconnected neural networks, 𝒳 and 𝒢. The network 𝒳 takes as input the time *t* and initial state *x*_0_, has a number of layers, and ends in an output layer predicting the state 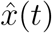. Automatic differentiation (not numerical differentiation, so exact) is employed in order to compute the time derivative 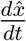, enabling direct evaluation of the governing ODEs. The network 𝒢 takes as input the predicted states 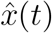 and, through a feedforward multilayer architecture, produces as its output an approximation 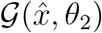 of the unknown dynamics *G*(*x*).

**Figure 1.**
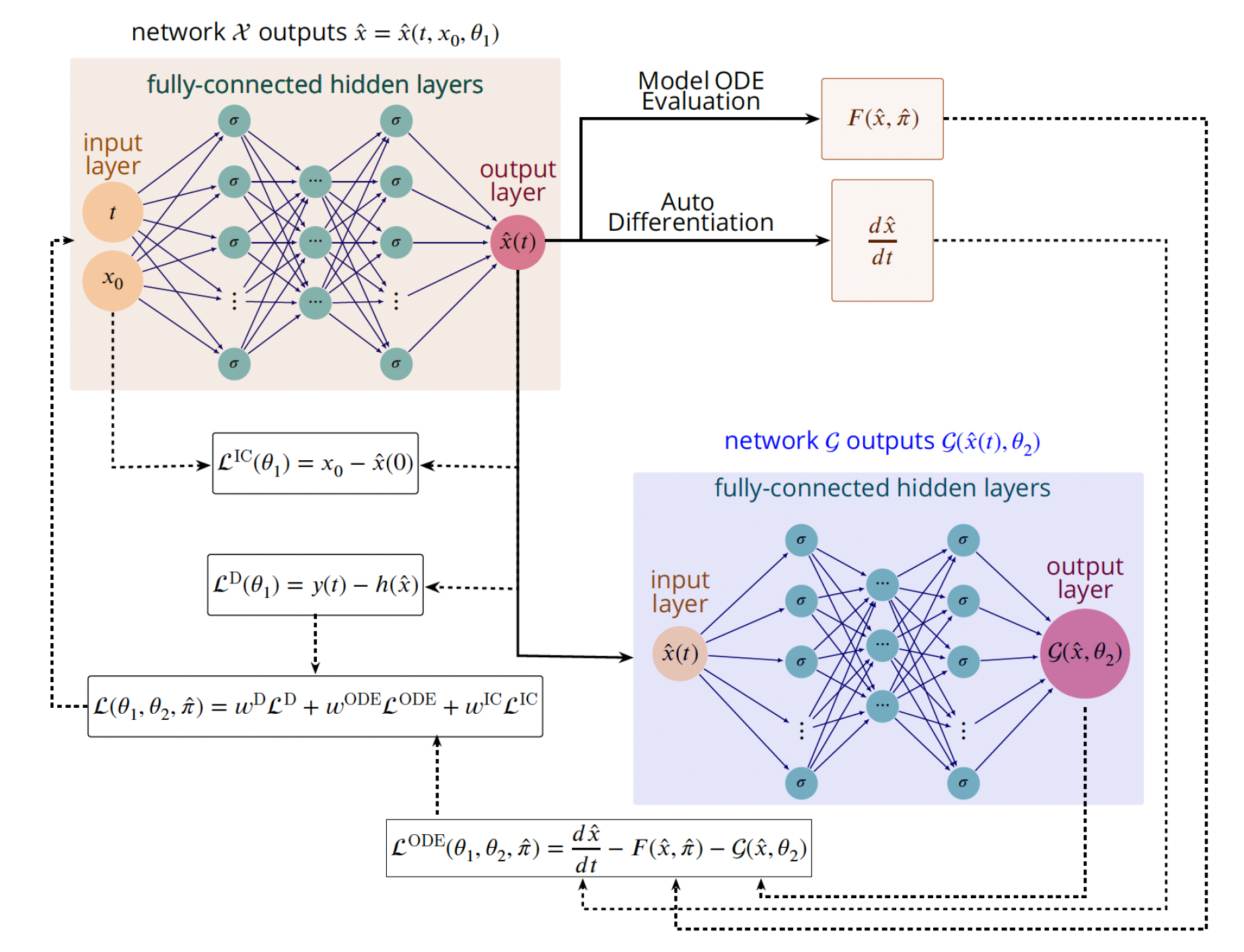
ePINNs architecture. ePINNs structure is composed of two embedded neural networks. The first one, 𝒳, has weights *θ*_1_, postulates a trajectory 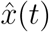 as a function of time and initial conditions *x*(0). The second one, 𝒢, has weights *θ*_2_ and takes as its input the postulated trajectory 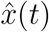 produced by 𝒳, producing a “correction term” 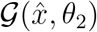 representing totally unknown dynamics. For simplicity, norms and sums over training data are not shown for the various loss functions *ℒ*; precise definitions are given in the text.

#### 2.1.1. Overall loss function structure

The two networks do not interact directly, but they both contribute to the ODE loss function 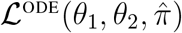, which, together with a penalty on the mismatch between 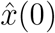 and *x*_0_, will be minimized so as to ensure that 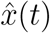 solves (2). To consistently train both neural networks using the governing ODEs and experimental data, we design a shared loss function for them, with the following components

1. *Data constraint, ℒ*^D^(*θ*_1_): Ensures that state predictions from 𝒳 match experimental observations.
2. *ODE constraint*,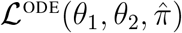: Ensures that derivative predictions 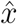, combined with unknown dynamics from, satisfy the ODE system.
3. *Initial condition constraint, ℒ*^IC^(*θ*_1_): Ensures that the initial state predictions 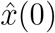 match the true initial conditions of all states (both observed and unobserved). This loss term is particularly crucial during the training process with partially observed dynamics, as the solution space of the known ODEs would otherwise span the entire R^*n*^ space, unless purposefully guided towards the true initial condition.

During training, we are given data *y*_*i*_ = *y*(*t*_*i*_) measured at a finite number of time points *t* = (*t*_1_, …, *t*_*N*_). For numerical experiments to evaluate the framework, we use a “ground truth” system of the form (1) to generate *y*(*t*), and we wish to estimate *π* (denoted by 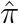) as well as find a network 𝒢 that approximates *G*. We assume that *x*_0_ is known, which is a realistic assumption in many problems, such as our case study below. The data loss term will be checked at the sampling times *t*_*i*_. We found in practice that it is best to employ an adaptive time sampling frequency which is higher during fast transition periods and is reduced during steady states, enabling effective separation of fast and slow dynamics.

We introduce coefficients, denoted by *w*^D^, *w*^ODE^, and *w*^IC^, respectively, to balance the relative contributions of *ℒ*^D^, *ℒ*^ODE^, and *ℒ*^IC^ into an overall loss function:

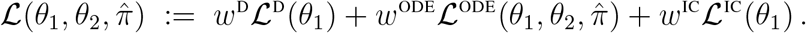

The explicit forms of losses are defined as follows:

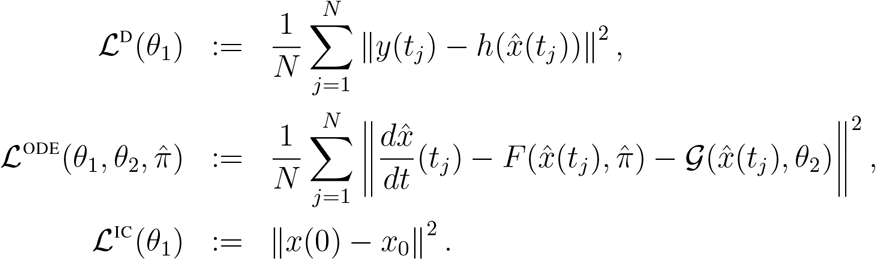

The loss functions are also shown in Figure 1 The loss components *ℒ*^D^(*θ*_1_) and *ℒ*^IC^(*θ*_1_) deal with mismatches between model predictions and observations *y*. On the other hand, the component 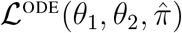 is not directly dependent on data and only relies upon ODE evaluations and automatic differentiation (AD), which computes 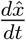 [24, 25]. AD calculates the gradients of the neural network outputs with respect to their inputs, ensuring accurate and efficient derivative estimation without explicit numerical differentiation.

Training is conducted using gradient-based optimization methods, such as the Adam optimizer [26], to update the parameters *θ*_1_, *θ*_2_, and 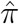 simultaneously. The optimized parameters 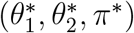 are obtained by minimizing the shared loss function 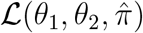, as follows:

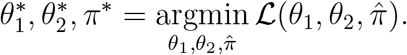

This simultaneous optimization ensures that the known dynamics, unknown dynamics, and system parameters are cohesively learned during the training process.

### 2.2. Case study: a resource competition model in synthetic biology

As described in the introductory Section 1.3, this work was motivated by the fact that engineered genetic circuits may not perform as expected when inserted into living cells, due to resource sharing.

In systems biology, one considers models in which internal states represent the concentrations of promoters, mRNAs, proteins, and other cellular players. Suppose that we are interested in a synthetic circuit in which a particular gene ultimately leads, through transcription, translation, and post-translational modifications, to the expression of a desired “output” protein, which we will denote by *Y*. In applications of synthetic biology, *Y* is a protein to be manufactured by a cell. For example, insulin is such a protein, in fact a small peptide hormone. Recombinant insulin is a synthetic form of human insulin that is used to treat diabetes mellitus by regulating blood sugar levels. Recombinant insulin is made by inserting the human insulin gene into genetically engineered bacteria (*E. coli*) or yeast (*S. cerevisiae*). These microorganisms produce insulin that is harvested and purified for pharmaceutical use.

We view the total concentration of the input gene as an “input” to the gene expression component, which we denote by *U* and assume is constant. Ideally, after the system settles to a steady state, the output concentration of *Y* will stabilize to a desired value. The process that leads from *U* into *Y* is shown in the green box in Figure 2.

**Figure 2.**
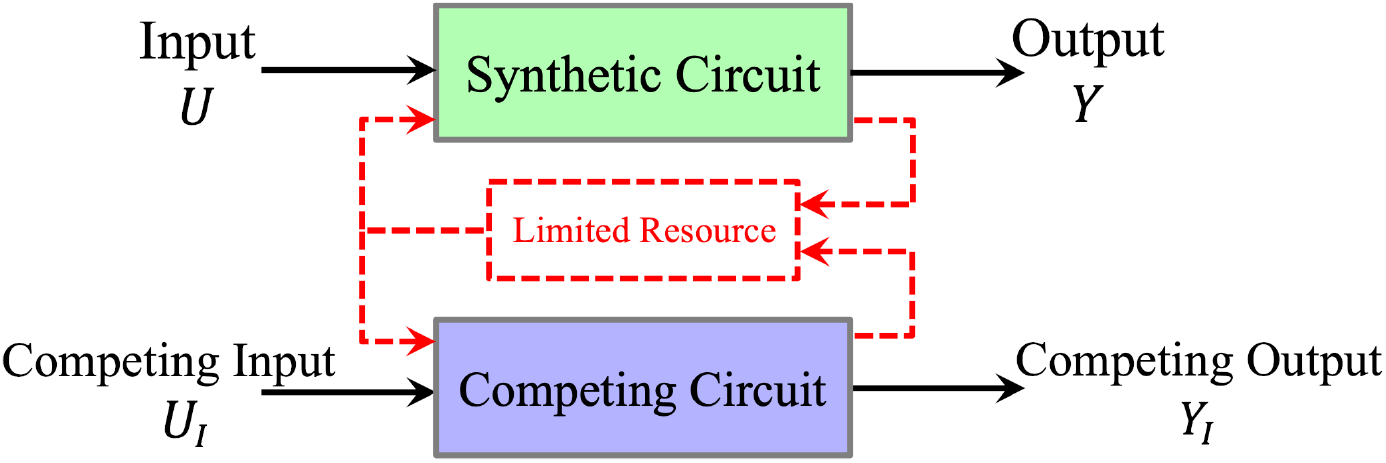
A synthetic circuit with one competing circuit.

We prefer to think of *U* as the promoter associated to the given gene, because the promoter may be free or it might be bound to RNA polymerase (RNAP) molecules. We will denote the concentration of the unbound (free) promoter by *U*. RNAPs are the “machines” that transcribe into an intermediate component, mRNA, the genetic information encoded in DNA. Subsequently, mRNA molecules guide the production of the protein *Y*, a process that is facilitated by ribosomes, which are the “machines” that help translate mRNA information into aminoacid sequences. Such a gene transcription process typically shares the use of one or more limited resources (shown as a red box in the figure) which are also used by other “competing” (or “interfering”) genes (denoted by subscripts *I*). These cellular resources include notably ribosomes (*R* in the model to be described) and RNA polymerase (RNAP) (*Q*). For simplicity, let us lump all such competing circuits into one (blue box in the figure). We will subscripts *I* to denote components of this other circuit; it has input *U*_*I*_ and produces protein *Y*_*I*_.

As a convention, we will use italicized variables (such as *M*) to denote concentrations in differential equations, and use roman variables (such as M) to denote the corresponding chemical species. With this convention, a simple model of the chemical reactions associated with the two-gene competing system is defined as follows:

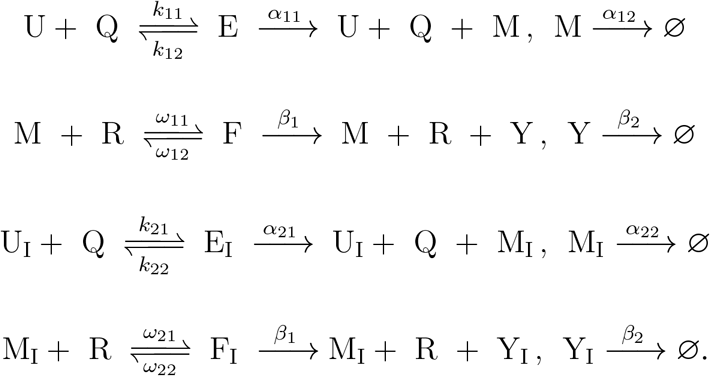

The first two sets of reactions describe the transcriptional and translational processes of the synthetic circuit, including degradation/dilution processes, while the last two sets describe the same processes for the competing circuit. There are five state variables for each circuit: *U, E, M, F, Y* for the synthetic circuit, and *U*_*I*_, *E*_*I*_, *M*_*I*_, *F*_*I*_, *Y*_*I*_ for the competing circuit. Additionally, there are two shared cellular resources, *R* and *Q*, resulting in a 12-dimensional full system. The state variables corresponding to each species are summarized in Table 1.

**TABLE 1.** State variables for model with two circuits sharing RNAP and ribosomes.

| Synthetic Circuit | Competing Circuit | Resources |
| --- | --- | --- |
| $U$: free promoter | $U_I$: free promoter | $R$: ribosome |
| $E$: promoter-RNAP complex | $E_I$: promoter-RNAP complex | $Q$: RNAP |
| $M$: mRNA | $M_I$: mRNA | |
| $F$: mRNA-ribosome complex | $F_I$: mRNA-ribosome complex | |
| $Y$: output protein | $Y_I$: output protein | |

A mass action kinetics representation of this system, including kinetic constants, is as follows.

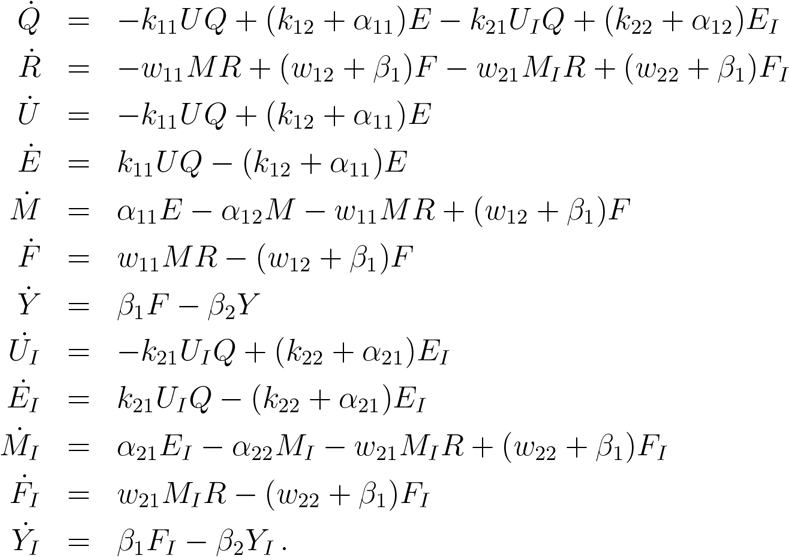

It is easy to see from the equations that the following concentrations remain constant over time:

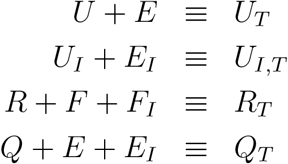

(for example, the time derivative of *U* (*t*) + *E*(*t*) is zero) so these represent conservation laws that allow one to reduce the equations (given a particular set of initial conditions) to an 8-dimensional system. The subscripts “*T* “stand for “total.” Natural initial conditions are *Q*(0) = *Q*_*T*_, *R*(0) = *R*_*T*_, *U* (0) = *U*_*T*_, *U*_*I*_(0) = *U*_*I,T*_, with the remaining variables set to zero. Using these conservation laws, we may eliminate variables as follows:

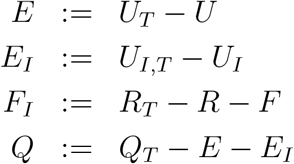

and we obtain the following set of equations, which we will call the “8D” model:

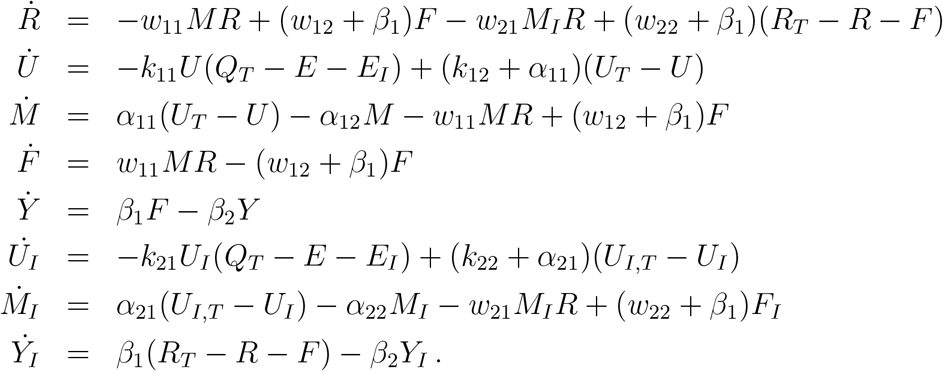

RNAP and ribosome competition give rise to different behaviors, studied for example in [27]. Let us focus on the most common scenario of limited ribosome availability but abundant RNAP. In this case, we assume that *R* is the bottleneck resource, while *Q* is maintained at a constant value *Q*(*t*) ≡ *Q*_*T*_. As a simplified model in that case, in the 12D system we use only the first there conservation laws, and we drop the differential equation for *Q*(*t*), replacing *Q* by *Q*_*T*_ in all the equations. This results in a “abundant-*Q*_*T*_ 8D” model (not shown). Next note that *Y*_*I*_ does not affect the first 7 equations. Moreover, in our application, we will assume that *Y*_*I*_ cannot be measured, as it represents protein produced by unknown genes. The model obtained by dropping the *Y*_*I*_ equation from the abundant-*Q*_*T*_ 8D model will be called the “7D” system:

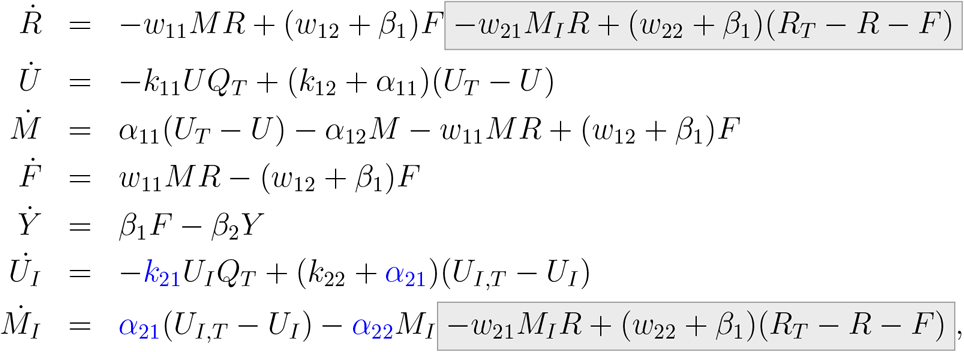

with initial condition (*R*_0_, *U*_0_, *M*_0_, *F*_0_, *Y*_0_, *U*_*I*0_, *M*_*I*0_)^⊤^ = (*R*_*T*_, *U*_*T*_, 0, 0, *U*_*I,T*_, 0)^⊤^. We consider the kinetic rates *π* = (*α*_21_, *α*_22_, *k*_21_) (marked in blue for emphasis) as unknown model parameters, while the remaining parameters are known. There is also a “black box” competition effect (grayed-out) that impacts the amount of free ribosomes (*R*) available to *M*. This is the *G* part of the system that will be approximated by the neural network 𝒢. The vector field “*F* (*x, π*)” corresponds to the non-grayed-out part of the equations. (Not to be confused with *F* also denoting one of the components of the state *x* = (*R, U, M, F, Y, U*_*I*_, *M*_*I*_)^⊤^.) We will take the output as (*Y, M, M*_*I*_). Using *Y* instead of *F* is more realistic (as this will typically represent a reporter protein, for example) than using the complex *F* as an output. We assume that we can individually measure *M* and *M*_*I*_. Such a measurement could be obtained by labeling the target mRNA *M* and separately measuring the total amount of transcripts in the cell, so that *M*_*I*_ would be the difference between these two numbers. The structural identifiability analysis of the unknown model parameters under this set of outputs is presented in 2.3.1.

Of course, if we knew that the interaction between *M*_*I*_ and *R* has this particular form in the gray box, the problem would be reduced to the parameter-fitting problem of finding *w*_21_, *w*_22_, and *β*_1_ in addition to *α*_21_, *α*_22_, and *k*_21_. However, we want to allow for the possibility of more general and unknown interactions. These might include ribosome stalling, use of rare codons, proofreading steps, ribosomal frameshifting, ribosome pausing and translational regulation, and other processes. Generally, a multi-step translational model may be more appropriate, such as the ribosome flow model [28] and in particular the splitting of the ribosome pool by use of orthogonal ribosomes [29].

A final possible simplification is as follows. Under an assumption of rapid binding and unbinding of RNAP to promoters, we assume an equilibrium value for the free promoters *U* and *U*_*I*_, and absorb these into the rates of transcription *α*_11_ and *α*_21_ respectively. Thus we arrive to the following “5D” system:

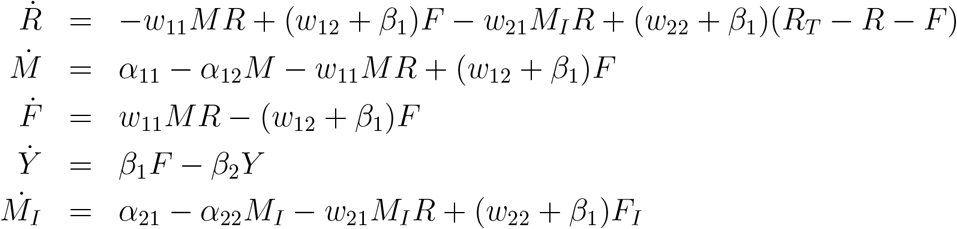

The protein *Y* is typically measured in experiments, so we did not drop the equation for *Y* in the 7D system. However, in order to study a minimal model, we will drop it from the equations in the 5D model, resulting in a “4D” model to be studied:

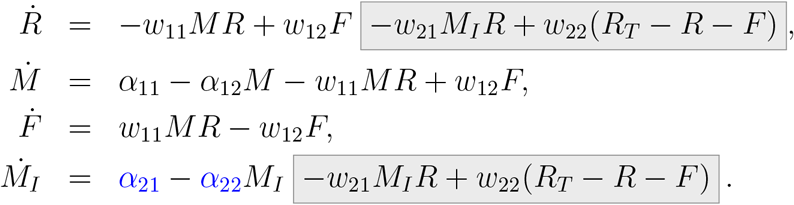

(We have absorbed *β*_1_ into *w*_12_ and *w*_12_.) The variables are *R* (free ribosomes), *M* and *M*_*I*_ (target and competing mRNA), and *F* (ribosomes bound to *M*, thought of as production rate of *Y*). The initial condition is (*R*_0_, *M*_0_, *F*_0_, *M*_*I*0_)^⊤^ = (*R*_*T*_, 0, 0, 0)^⊤^. We view this system as consisting of a partially unknown competing subsystem represented by the *M*_*I*_ variable, in which the unknown parameters are the two kinetic rates *π* = (*α*_21_, *α*_22_) (marked in blue for emphasis). We will use this 4D system in order to numerically explore how partial mechanistic knowledge can improve data fits using ePINNs.

One last issue in setting up the model for numerical experimentation is the choice of output *y*. Observe from the equations that even if all three of *R, M, F* were measured, it would be impossible to get any information whatsoever about *M*_*I*_, since *M*_*I*_ only affects *R* (and indirectly *M* and *F*) only through the totally unknown black box *G*(*x*). This means that there is no hope of identifying the unknown parameters. Therefore, to make the problem nontrivial, we have picked the output *y* = (*F, M* + *M*_*I*_)^⊤^. In other words, we assume that we can measure the expressed protein of interest as well as the total amount of mRNAs. The structural identifiability analysis of the unknown model parameters under this set of outputs is presented in 2.3.2.

By focusing on the reduced 7D and 4D systems, we aim to highlight the key concepts more clearly while avoiding unnecessary complexity.

### 2.3. Structural Observability and Parameter Identifiability Analysis

To confirm the feasibility of the hybrid state reconstruction, parameter identification, and hidden mechanism estimation problem—that is, whether ePINNs can successfully recover all states, unknown parameters, and hidden dynamics under the specified output configurations and unknowns for both the 7D and 4D models—we perform a structural observability and identifiability analysis using the STRIKE-GOLDD toolbox [30]. This toolbox is based on the methodology introduced in [31], which applies the extended Observability Rank Condition (extended ORC) for systems with unmeasured inputs. In this approach, the system is augmented by incorporating unknown parameters and unmeasured inputs (along with their time derivatives up to a finite order) into an extended state vector. Lie derivatives of the output functions are then recursively computed with respect to this augmented system, and the resulting observability-identifiability matrix is constructed from the Jacobians of these Lie derivatives.

To address the computational complexity of symbolic differentiation, STRIKE-GOLDD uses a power series-based semi-numerical implementation that relies on Newton iteration and modular arithmetic. This enables efficient rank computation of the observability-identifiability matrix, even for systems with rational nonlinearities and unmeasured inputs, by evaluating the matrix over a finite field. A system is deemed structurally observable if this matrix attains full rank.

Furthermore, we perform analytical parameter identifiability analysis for each model under its respective output configuration, assuming the unknown dynamics are reconstructible (as verified by STRIKE-GOLDD). This serves to confirm that the chosen set of unknown parameters is structurally identifiable and therefore feasible to infer.

#### 2.3.1. 7D Model Analysis

In this section, we study the structural observability and identifiability of the 7D model. For the purpose of this analysis, we consider a broader scenario in which the parameter *k*_22_ is also treated as unknown. This allows us to gain insight into how the choice of problem formulation affects parameter identification when using ePINNs. To investigate the structural identifiability of the model parameters *k*_21_, *k*_22_, *α*_21_, and *α*_22_, we examine a simplified subsystem (presented below) in which only the trajectory of *M*_*I*_(*t*) is observed, while *U*_*I*_(*t*) remains unmeasured. For this analysis, we exclude any black-box (unknown) dynamics, focusing solely on the parametric component of the model. The governing dynamics will then be:

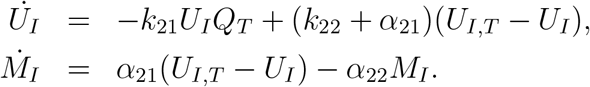

We define the state vector and output as

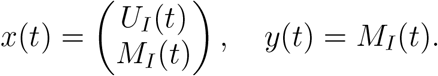

This system can be written in state-space form:

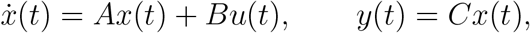

where *u*(*t*) ≡ 1 and

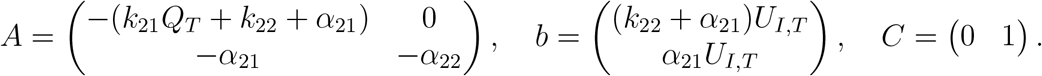

The first and second derivatives of the output *y*(*t*) are given by:

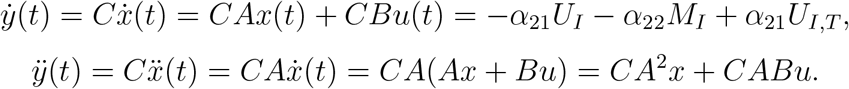

Since *A* and *B* are constant, 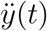 is a linear combination of the state variables *U*_*I*_ and *M*_*I*_ with coefficients involving the model parameters. Substituting back, we can express 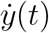 and 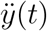 in terms of *y*(*t*) and its derivatives:

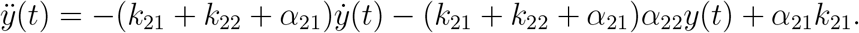

This implies that only specific combinations—*α*_21_*k*_21_, *k*_21_ + *k*_22_ + *α*_21_, and *α*_22_—appear in the output equation. Hence, *α*_22_ is structurally identifiable, while the individual parameters *k*_21_, *k*_22_, and *α*_21_ are not separately identifiable from *M*_*I*_(*t*) alone.

It is easy to show that if any one of the parameters *k*_21_, *k*_22_, or *α*_21_ is known, the remaining parameters become structurally identifiable. This result is fully consistent with the structural identifiability analysis performed using STRIKE-GOLDD, which confirms that the indeterminacy lies in the combinations of these three parameters when *M*_*I*_(*t*) is the only observed output. Therefore, to keep the identifiability problem feasible for ePINNs, we assume that *k*_22_ is known.

#### 2.3.2. 4D Model Analysis

Here, we investigate further the observability and identifiability of the 4D model described earlier. We assume the observed outputs are:

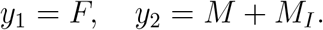

The first and second derivatives of the outputs will then be:

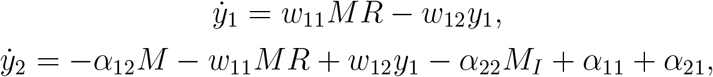

and the second derivative is given by:

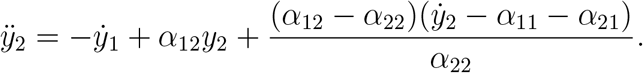

From this form, it is evident that both *α*_21_ and *α*_22_ are structurally identifiable.

We also applied the STRIKE-GOLDD analysis to the 4D model comprising the state variables *R, M, F*, and *M*_*I*_, with two unknown parameters, *α*_21_ and *α*_22_. The unmodeled black-box mechanism was modeled as an unknown, time-varying input. The analysis showed that when the outputs were limited to the protein production rate *F* and the total mRNA concentration *M* + *M*_*I*_, the observability-identifiability matrix achieved full rank. This indicates that all four state variables, as well as both unknown parameters, are locally structurally observable and identifiable, which supports our analytical identifiability result. Thus, in theory, this minimal measurement configuration suffices to reconstruct the full system dynamics and parameter values, despite the presence of an unmeasured input.

### 2.4. Partially uncertain model structures

As discussed in the introductory Section 1.2, in order for the problem to be well-posed, we need to impose some *structure* to the uncertainty. In order to do this, we will partition the states into two components:

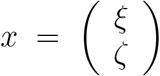

where *ξ* ∈ ℝ^*p*^, *ζ* ∈ ℝ ^*q*^, and *p*+*q* = *n*. The first block of variables will collect those mechanistic parts for which parameters are known.

We start with an example to illustrate the general idea:

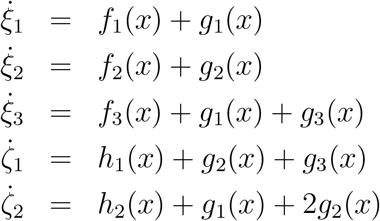

in which *p* = 3 and *q* = 2. We use lower case *ξ*_*i*_, *ζ*_*i*_ for the coordinates of *ξ, ζ*. Here *G* is the vector with coordinates *g*_1_, *g*_2_, *g*_1_ + *g*_3_, *g*_2_ + *g*_3_, and *g*_1_ + 2*g*_2_ that represents how three “neural networks” affect the dynamics, and *F* has the coordinates *f*_1_, *f*_2_, *f*_3_, *h*_1_, *h*_2_, where

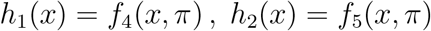

are the components of the mechanistic part of the model which contain the unknown parameters. As our focus is not on the values of the parameters *π*, we simply think of the *h*_*i*_’s as unknown functions that we wish to identify. Now, suppose that we have already obtained a model in which the coordinates of *x*(*t*), that is, the functions *ξ*_*i*_(*t*) and *ζ*_*i*_(*t*), have been found to fit given experimental data. There may be many sets of such functions *x*, obtained from different initial conditions or unmodeled factors. Since we know the functions *ξ*_*i*_(*t*), we may compute, at least in an ideal mathematical sense, their derivatives, and thus, subtracting the (known) function *f*_*i*_(*x*(*t*)), we also know the functions *γ*_1_(*t*) = *g*_1_(*x*(*t*)) and *γ*_2_(*t*) = *g*_2_(*x*(*t*)). Similarly, from the knowledge of *ζ*_1_(*t*) and *ζ*_2_(*t*), we know 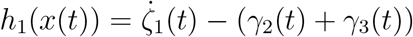and 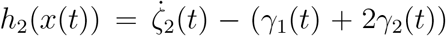. In summary, at least if the trajectories *x* are rich enough to explore the state space, we found the functions *h*_1_(*x*) and *h*_2_(*x*), which was our objective, We next make this discussion mathematically precise. We start by thinking of this example as follows. Let us introduce the following three matrices:

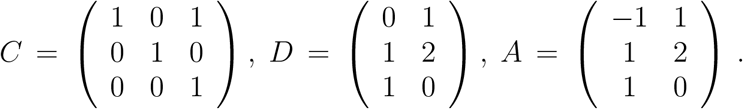

Observe that *D* = *CA*. This property explains why we could recover the functions *h*_*i*_, as we will discuss next.

**Definition**. A *Partially Uncertain Model Structure* **(PUMS)** is a system structure as follows:

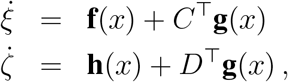

in which *x* = (*ξ, ζ*)^⊤^. The functions **f, h**, and **g** map R^*n*^ into R^*p*^, R^*q*^, and R^*r*^ respectively, and *C* ∈ ℝ^*r*×*p*^, *D* ∈ ℝ^*r*×*q*^.

A PUMS structure is *identifiable* if from knowledge of *x*(*t*) can recover **h** (i.e., parameters). More formally, if the following property holds:

*for all functions* ***g***, 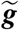, ***h***, *and* 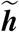, *if:*

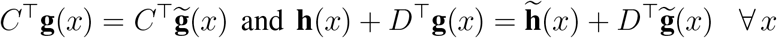

*then:*

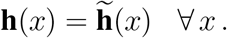

Note that our example can be written as a PUMS, with **g** = (*g*_1_, *g*_2_, *g*_3_)^⊤^.

The interpretation of the identifiability definition is as follows. If we know that **f, g**, and **h** and **f**, 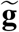, and 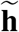, give the same trajectories 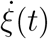 and 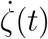, then necessarily the parameterized mechanistic parts **h** and 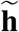 must have been the same. In Theorem 1 (Section 3.3) we provide a necessary and sufficient condition. Intuitively the condition will say that from *ξ* we can obtain *C*^⊤^**g**, because **f**(*x*) is known, and the condition given there then implies that *D*^⊤^**g** is known, so that **h** can be found from knowledge of this term as well as *ζ*.

## 3. Results

In this section, we explore our case study to demonstrate the implementation of the ePINNs approach for learning hidden mechanisms and predicting causal connections in the synthetic biology resource competition problem, and we provide a statement and proof of an identifiability result for PUMS.

We aim to explore the following research questions through this case study:

1. *System observability*: Using only the selected output measurements, can the trajectories of all states be accurately predicted?
2. *Parameter identification*: Can unknown parameters (chemical reaction rates, in this example) be inferred effectively?
3. *Missing dynamics prediction*: Can the “black-box” hidden mechanism (in this example, related to resource competition) be uncovered?
4. *Comparing scenarios:* Does even partial knowledge improve fits, compared to a purely “black box” approach?

We will evaluate several scenarios with varying levels of system knowledge for the described case study.

### 3.1. 4D Model

In this section, we use the 4D model and the parameters shown in Table 2 to generate the synthetic data used for training ePINNs. To simulate experimental uncertainty, we introduced additive Gaussian noise to the synthetic data. For each observed state variable *x*, the noise term *η*(*t*) was sampled from a normal distribution 𝒩 (0, *σ*^2^), where *σ* was set to 5% of the variability of *x* over the sampling period, denoted by Var(*x*). The resulting noisy state variables are given by 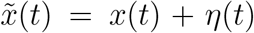, where *η*(*t*) ∼ 𝒩 (0, (0.05 ·Var(*x*))^2^). This ensures that the noise magnitude is proportional to the natural variability of each observed state over time. The noisy state trajectories 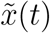 were used for training.

**TABLE 2.** Parameter values for the reduced 4D model used in simulations. Units are as appropriate (concentrations/second, etc), but these are arbitrary parameters picked only in order to illustrate results.

| Parameter | $\omega_{11}$ | $\omega_{21}$ | $\alpha_{11}$ | $\alpha_{21}$ | $\omega_{12}$ | $\omega_{22}$ | $\alpha_{12}$ | $\alpha_{22}$ | $R_T$ |
| --- | --- | --- | --- | --- | --- | --- | --- | --- | --- |
| Value | 1 | 1 | 1 | 3 | 3 | 7 | 3 | 5 | 20 |

We then evaluate the performance of the ePINNs approach for the defined resource competition problem with the same 4D dynamics under the following scenarios.

#### 3.1.1. Impact of full mechanistic knowledge on estimation and prediction with partial observations

To make the problem interesting (and more biologically realistic), we use only the partial observations represented by the output variable *y* = (*F, M* + *M*_*I*_)^⊤^ instead of the four state variables. One might reasonably ask whether the use of restricted information, meaning that we need to solve a simultaneous systems identification and observability problem, is the reason that some approaches might perform better than others. To control for this effect, we start by examining how the availability of full knowledge about system dynamics influences the estimation and prediction of behavior when only partial observations are available. In this case, the unknown mechanism and the blue parameters in our 4D system are assumed to be known. Our goal is to evaluate whether ePINNs can successfully reconstruct all state variables using only partial, noisy measurements (question (1)). This tests the method’s ability to infer unmeasured dynamics and assess observability. We assume that only data from total mRNA (*M* + *M*_*I*_) and the mRNA-ribosome complex (*F*) are available.

Figure 3 illustrates the results obtained from a “physically **un**informed” neural network (set the mechanistic loss weight *w*^ODE^ = 0), where predictions are represented by solid green lines and labeled as “NNs.” By comparison, Figure 4 shows the results from a physically informed neural network (*w*^ODE^ = 0.5), where predictions are represented by solid green lines and labeled as “ePINNs.” Both approaches provide accurate estimations for the observed data (*F* and *M* + *M*_*I*_).

**Figure 3.**
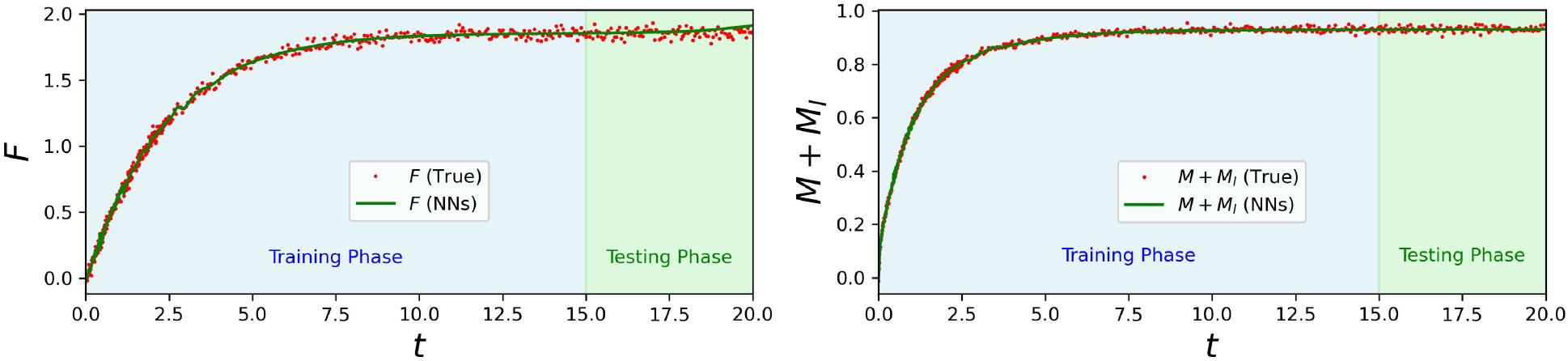
Physically uninformed neural network (*w*^ODE^ = 0) results for observables. Predictions for state variables *R, M* + *M*_*I*_ (solid green lines, labeled “NNs”) are shown using noisy data from *F* and *M* + *M*_*I*_ (red dots, labeled “True”). The physically uninformed network provides an accurate estimation for observed states.

**Figure 4.**
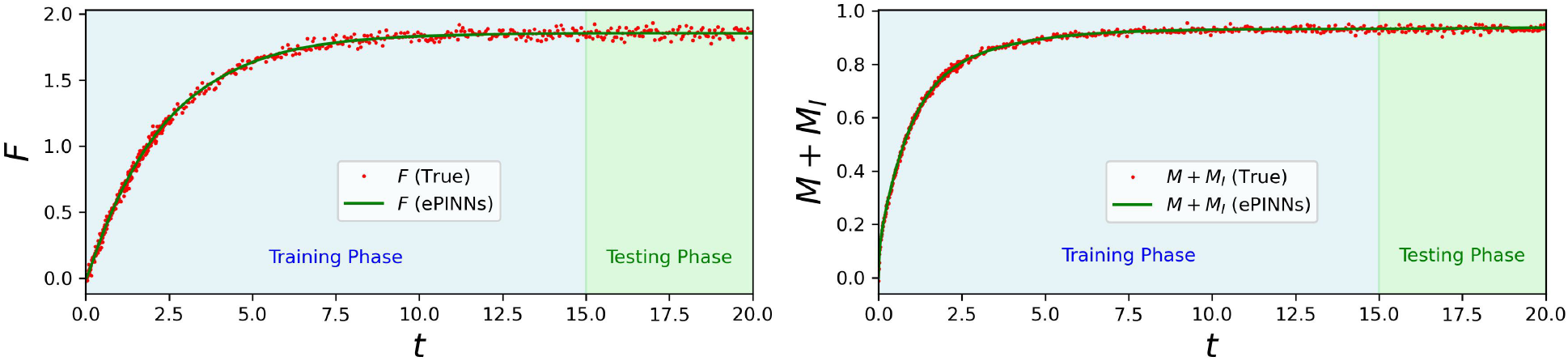
Physically informed neural network (*w*^ODE^ = 0.5) results for observables. Predictions for state variables *R* and *M* + *M*_*I*_ (solid green lines, labeled “ePINNs”) are shown using noisy data from *F* and *M* + *M*_*I*_ (red dots, labeled “True”). The physically informed network provides an accurate estimation for observed states.

Although both approaches utilize the available data, it is evident from Figures 5 and 6 that the physically uninformed neural network provides inaccurate estimations for the state components *R, M*, and *M*_*I*_, for which no direct data is available. However, as shown in Figure 6, the physically informed network effectively leverages the system’s physics knowledge to achieve accurate estimations for all states, even in the absence of direct measurements of all states (question (4)).

**Figure 5.**
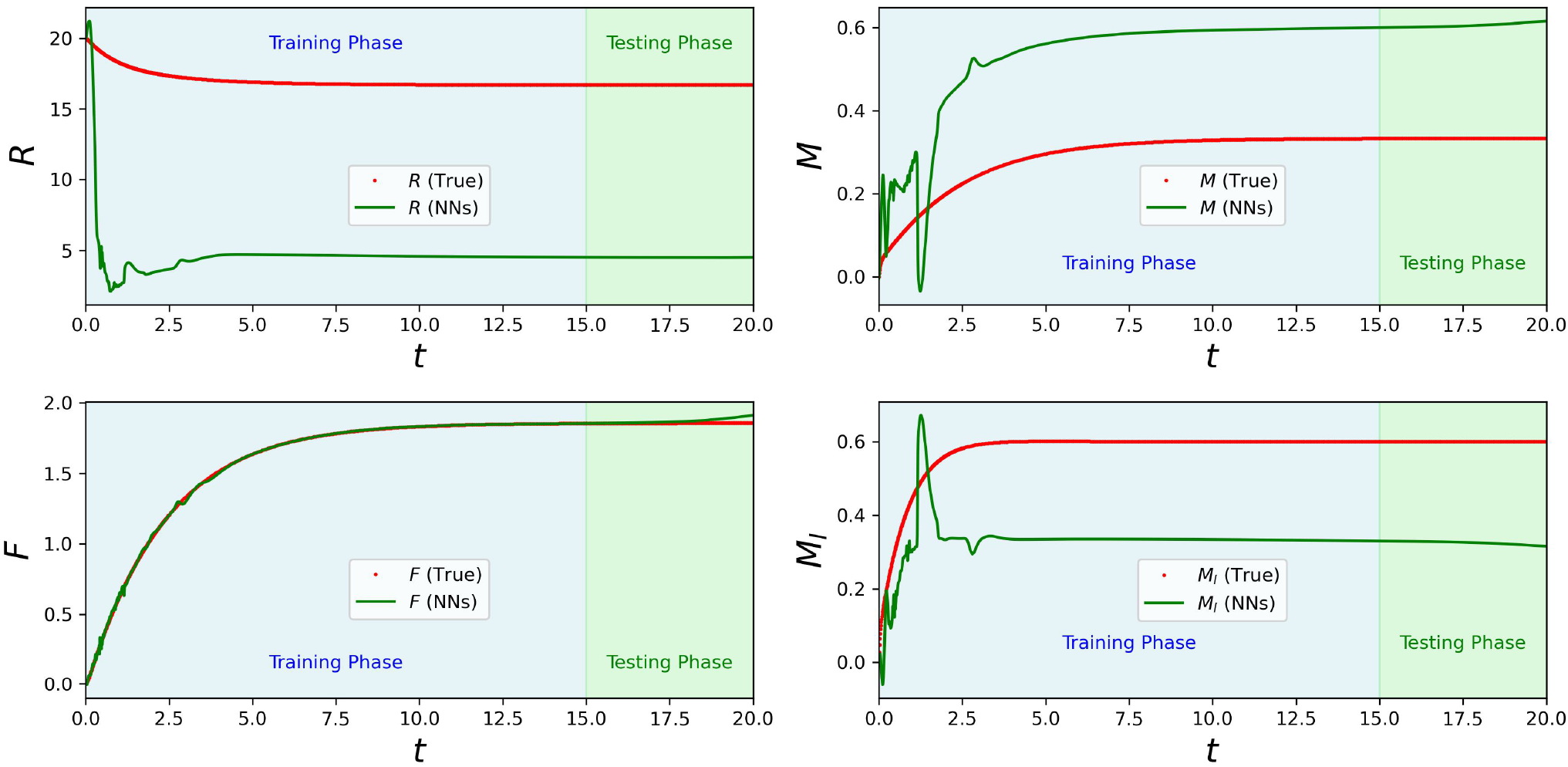
Physically uninformed neural network (*w*^ODE^ = 0) predictions for all states. Predictions for state variables *R, M*, and *M*_*I*_ (solid green lines, labeled “NNs”) are shown using noisy data from *F* and *M* + *M*_*I*_ (red dots, labeled “True”). The physically uninformed network fails to accurately estimate unobserved states due to the absence of physics knowledge.

**Figure 6.**
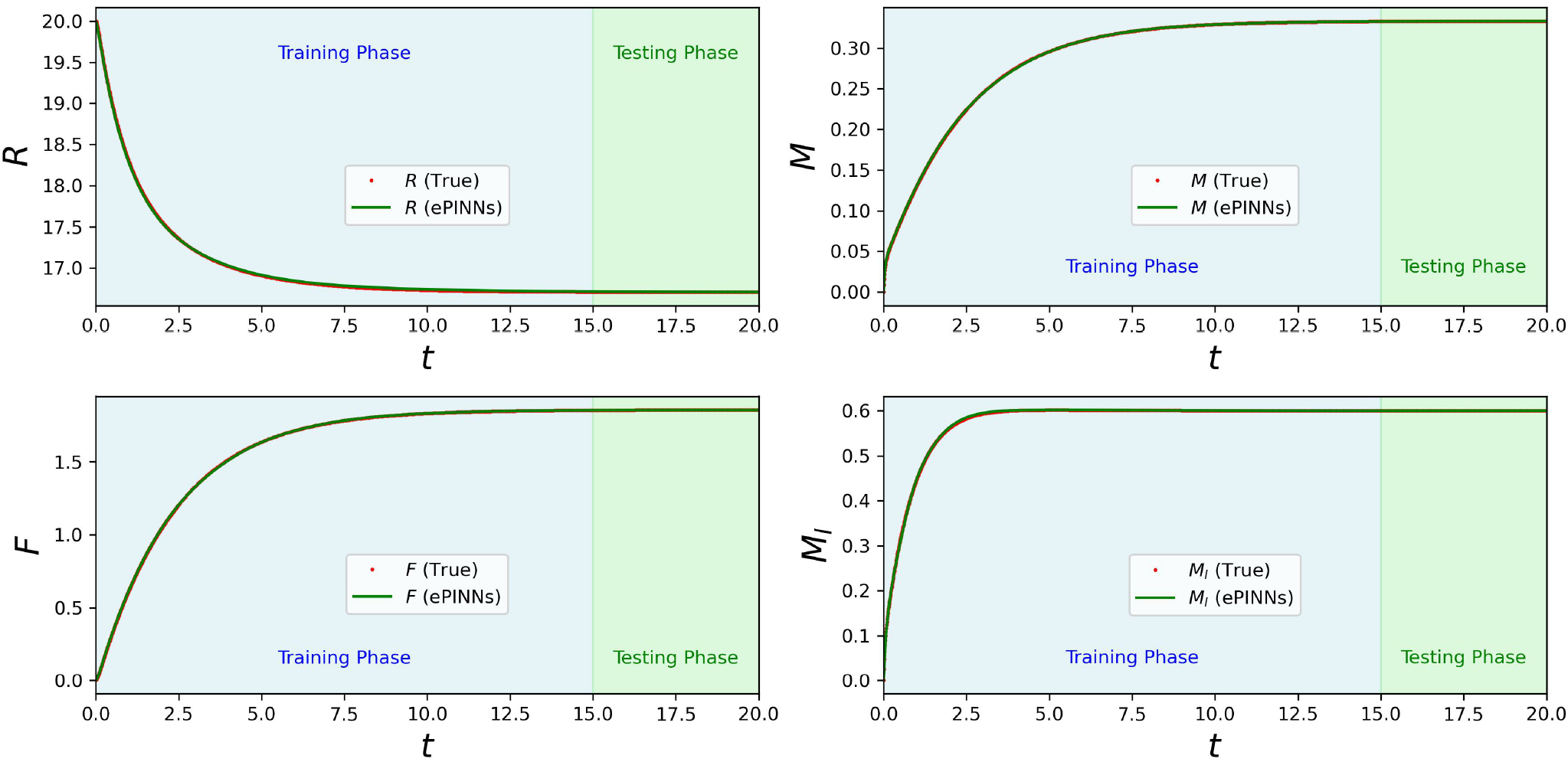
Physically informed neural network (*w*^ODE^ = 0.5) predictions for all states. Predictions for state variables *R, M*, and *M*_*I*_ (solid green lines, labeled “ePINNs”) are shown using noisy data from *F* and *M* +*M*_*I*_ (red dots, labeled “True”). The physically informed network accurately estimates all states, leveraging system dynamics even in the absence of direct data.

We note that both training processes (for the informed and uninformed cases discussed above) were conducted under identical conditions, network structures, and observed noisy data to ensure a fair comparison. In both cases, a learning rate of 0.01 was employed, with the first neural network consisting of 4 fully connected hidden layers, each containing 64 neurons. The “softplus” activation function was chosen for both scenarios. The training phase spanned from 0 to 15, while the testing phase extended from 15 to 20 to evaluate the neural networks’ accuracy in predicting state behaviors beyond the training period. An epoch count of 180*k* was selected uniformly across all cases.

#### 3.1.2. Partially observed and partially known system identification

This scenario represents the most realistic case, where both full state measurements and system knowledge are incomplete. Missing data and unknown interactions challenge the ePINNs framework to simultaneously infer the unknown parameters *α*_21_ and *α*_22_ (question (2)), as well as the missing terms in the gray box (question (3)).

Training data is assumed to be available only for the total mRNA (*M* + *M*_*I*_) and the mRNA-ribosome complex (*F*), leaving the system partially observed. The first neural network (𝒳) estimates the trajectory as a function of time, while the second neural network (𝒢) models the unknown ribosome competition term within the gray box in our 4D system. The trained model incorporates the dynamics of 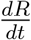 and 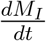 as follows:

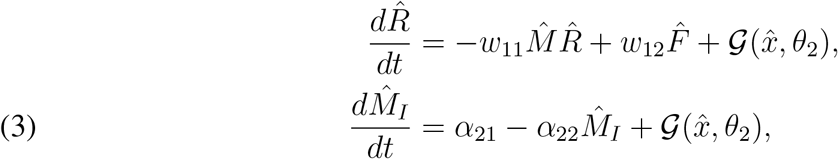

where 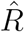 and 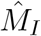 are the first neural network output states corresponding to *R* and *M*_*I*_, respectively.

The learning patterns for the unknown parameters *α*_21_ and *α*_22_ are presented in Figure 7, showing the convergence of ePINNs predictions (solid green lines) to their true values (dashed red lines) using noisy observations from *F* and *M* + *M*_*I*_.

**Figure 7.**
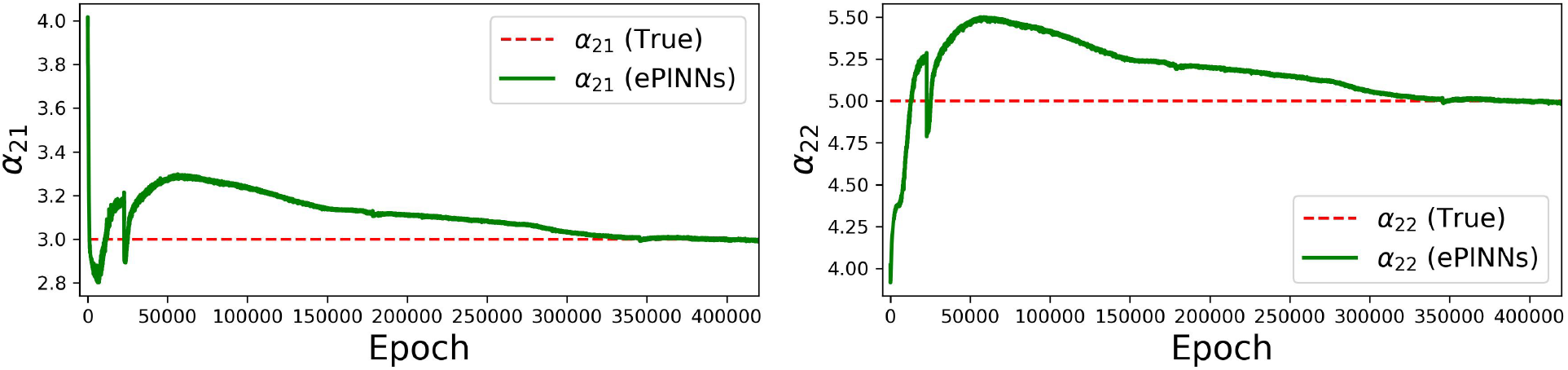
Learning patterns for unknown parameters *α*_21_ and *α*_22_ with partial data and missing dynamics. The solid green lines (labeled “ePINNs”) represent the estimated values, while the dashed red lines indicate the true values. The training process uses noisy observations from *F* and *M* + *M*_*I*_, demonstrating ePINNs’ ability to identify unknown parameters.

To evaluate missing dynamics prediction, Figure 8 illustrates how ePINNs reconstruct the hidden competition effect *G*(*x*). Despite only partial data availability, the model successfully learns the underlying mechanism, with the predicted dynamics (solid green line) closely matching the ground truth (red dots).

**Figure 8.**
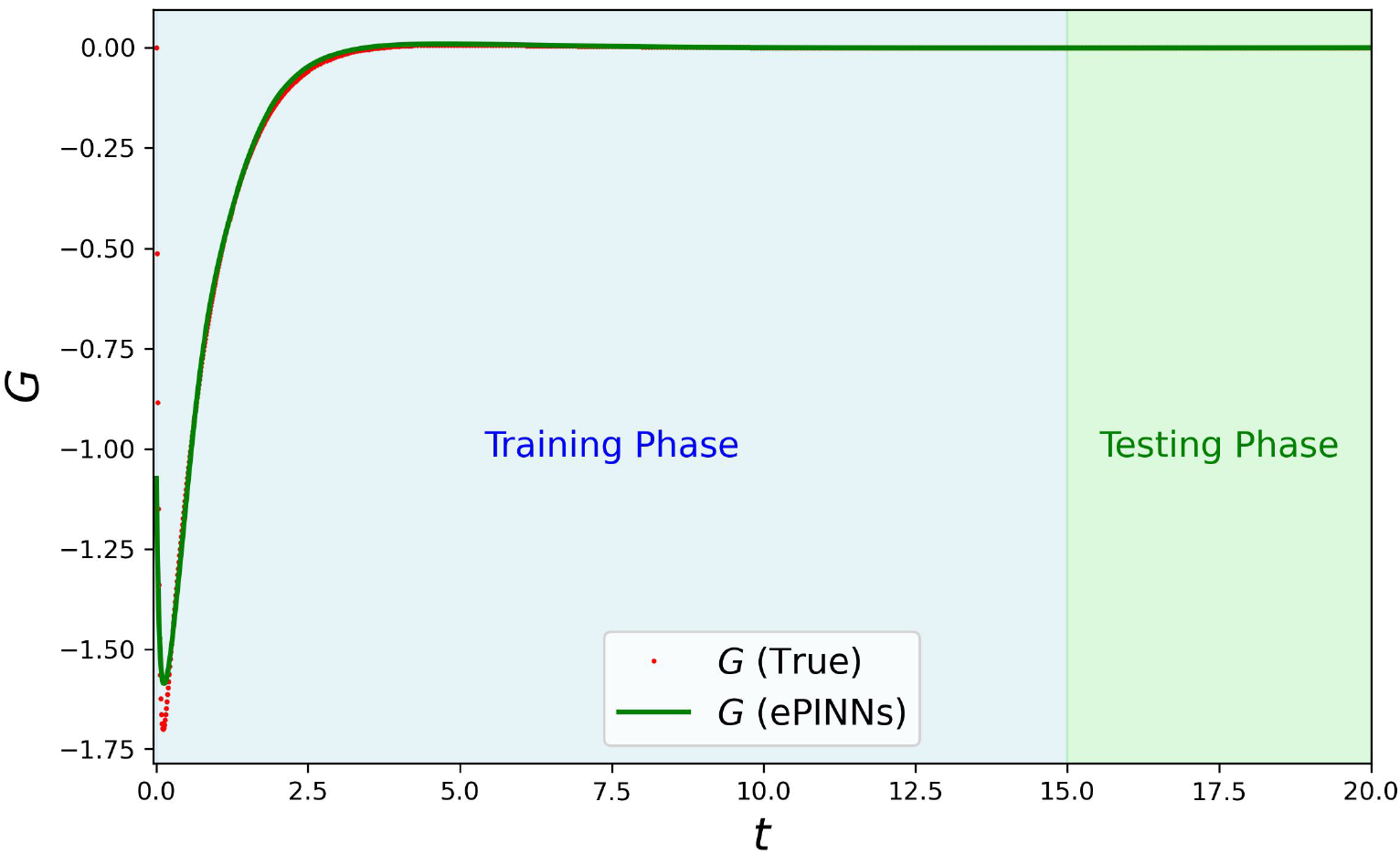
ePINNs prediction of missing dynamics *G*(*x*) with partial data. The solid green line (labeled “ePINNs”) represents the model’s prediction for the unknown dynamics, while red dots (labeled “True”) indicate ground truth values. The training uses noisy observations from *F* and *M* + *M*_*I*_, demonstrating the network’s ability to infer hidden system components.

Finally, Figure 9 provides a comprehensive evaluation of system observability, parameter identification, and missing dynamics prediction. The ePINNs model successfully reconstructs all state variables (*R, M, F, M*_*I*_) by leveraging available data and mechanistic constraints, highlighting its effectiveness in hybrid modeling scenarios. The training process for this example was conducted with a learning rate of 0.02. The first neural network consisted of 4 fully connected hidden layers, each containing 64 neurons, while the second neural network had 2 layers with 64 neurons each. The “softplus” activation function was used for both networks. An epoch count of 420k was applied uniformly across all cases.

**Figure 9.**
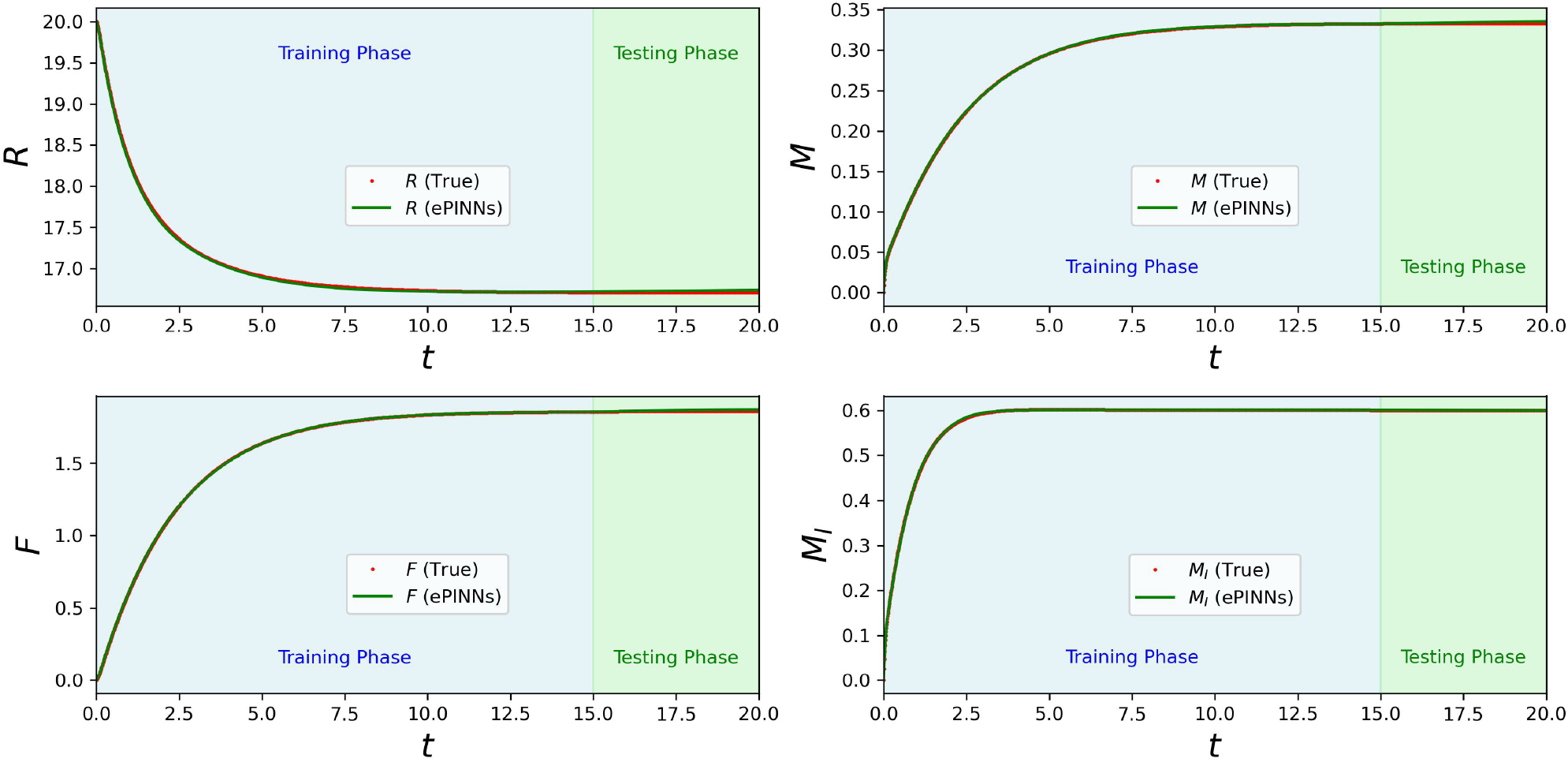
Predictions for all state variables with partial data and missing dynamics. The solid green lines (labeled “ePINNs”) show the predicted states *R, M, F*, and *M*_*I*_, while red dots (labeled “True”) represent the observed data for *F* and *M* + *M*_*I*_. The results illustrate the effectiveness of ePINNs in reconstructing unobserved states using physics-informed constraints.

### 3.2. 7D Model

In this section, we use the 7D model and the parameters shown in Table 3 to generate the synthetic data used for training ePINNs. To simulate experimental uncertainty, we use the same method explained for the 4D model. We use *Y, M*, and *M*_*I*_ as outputs for the training purposes.

**TABLE 3.** Parameter values for the 7D model used in simulations. Units are as appropriate (concentrations/second, etc), but these are arbitrary parameters picked only in order to illustrate results.

| Parameter | $\omega_{11}$ | $\omega_{21}$ | $\alpha_{11}$ | $\alpha_{21}$ | $k_{11}$ | $k_{21}$ | $\beta_1$ | $Q_T$ | $U_T$ |
| --- | --- | --- | --- | --- | --- | --- | --- | --- | --- |
| Value | 1 | 1 | 1 | 3 | 1 | 1 | 2 | 18 | 15 |
| Parameter | $\omega_{12}$ | $\omega_{22}$ | $\alpha_{12}$ | $\alpha_{22}$ | $k_{12}$ | $k_{22}$ | $\beta_2$ | $R_T$ | $U_{I,T}$ |
| Value | 3 | 7 | 3 | 5 | 8 | 12 | 1 | 20 | 26 |

For this example, training was performed with a scheduled learning rate. The architecture consisted of two neural networks: the first network featured four fully connected hidden layers with 64 neurons per layer, while the second comprised two layers of equal size. The “softplus” activation function was employed throughout. This model was trained for 280,000 epochs.

Figure 10 shows the evolution of the learned parameters *α*_21_, *α*_22_, and *k*_21_. The ePINNs estimates closely track the true values, indicating accurate parameter identification from limited observations.

**Figure 10.**
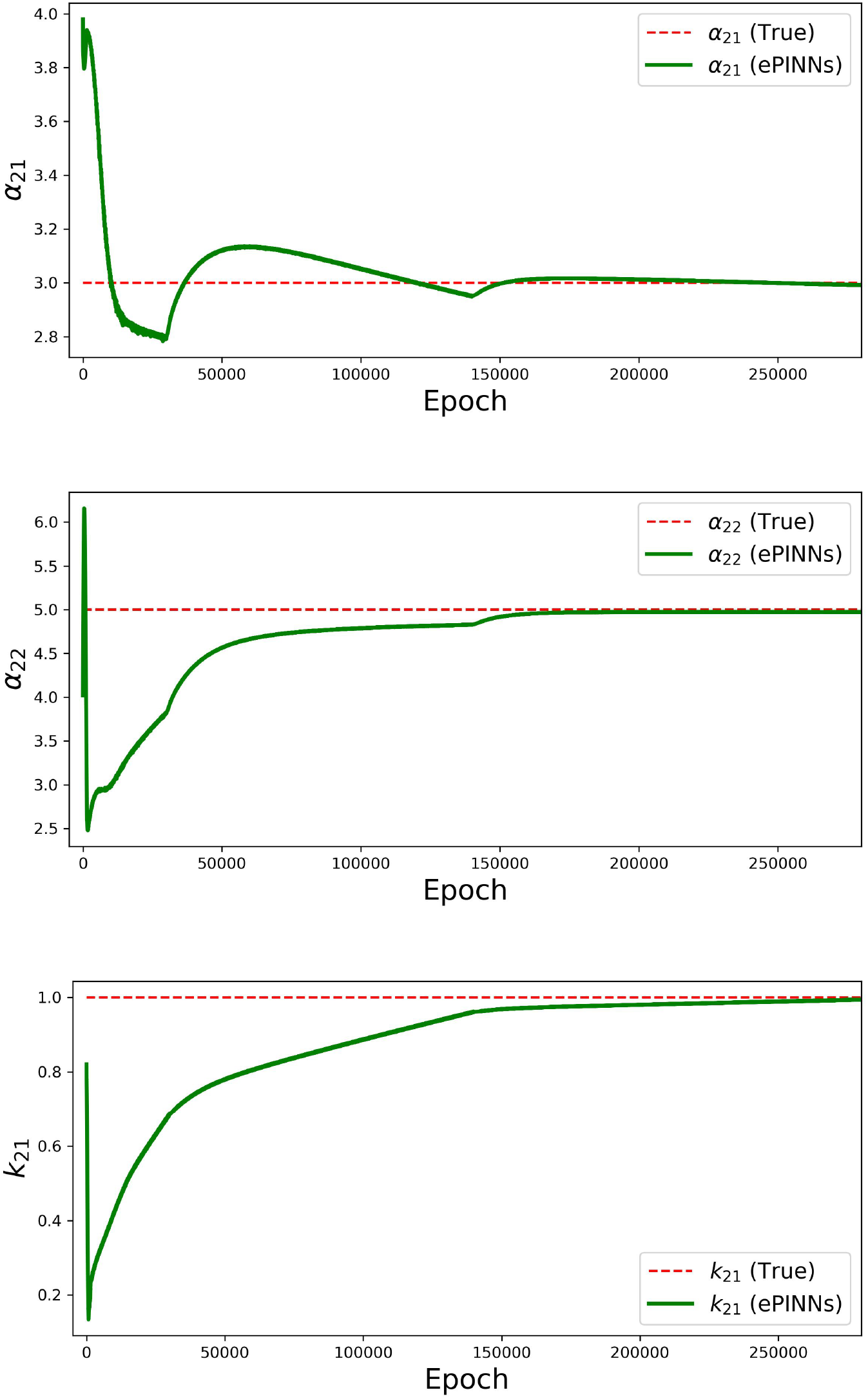
Learning patterns of unknown parameters *α*_21_, *α*_22_, and *k*_21_ using ePINNs. Solid green lines (labeled “ePINNs”) indicate ePINNs estimates; dashed red lines (labeled “True”) indicate true values.

Figure 11 presents the predictions for all state variables. Although only *Y, M*, and *M*_*I*_ were used for training, the model is able to recover the full state trajectories by leveraging the system dynamics.

**Figure 11.**
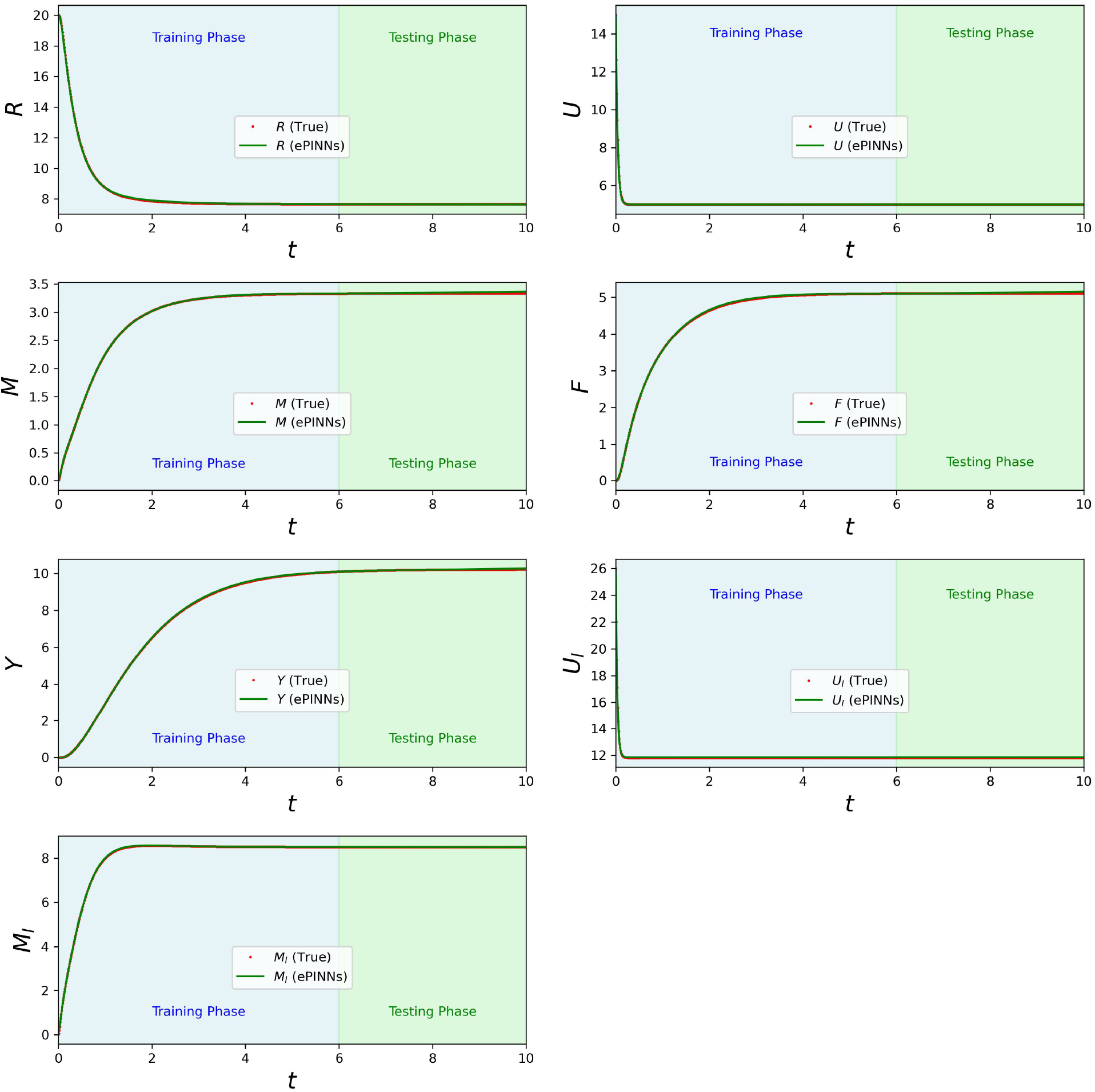
Predictions for all state variables of the 7D model with partial data and missing dynamics. The solid green lines (labeled “ePINNs”) show the predicted states *R, U, M, F, Y, U*_*I*_, and *M*_*I*_, while red dots (labeled “True”) represent the observed data for *Y, M*, and *M*_*I*_. The results illustrate the effectiveness of ePINNs in reconstructing unobserved states using physics-informed constraints.

Finally, Figure 12 demonstrates ePINNs’ ability to infer the unknown dynamics *G*(*x*). The predicted curve aligns well with the ground truth, despite the presence of noise and partial observations.

**Figure 12.**
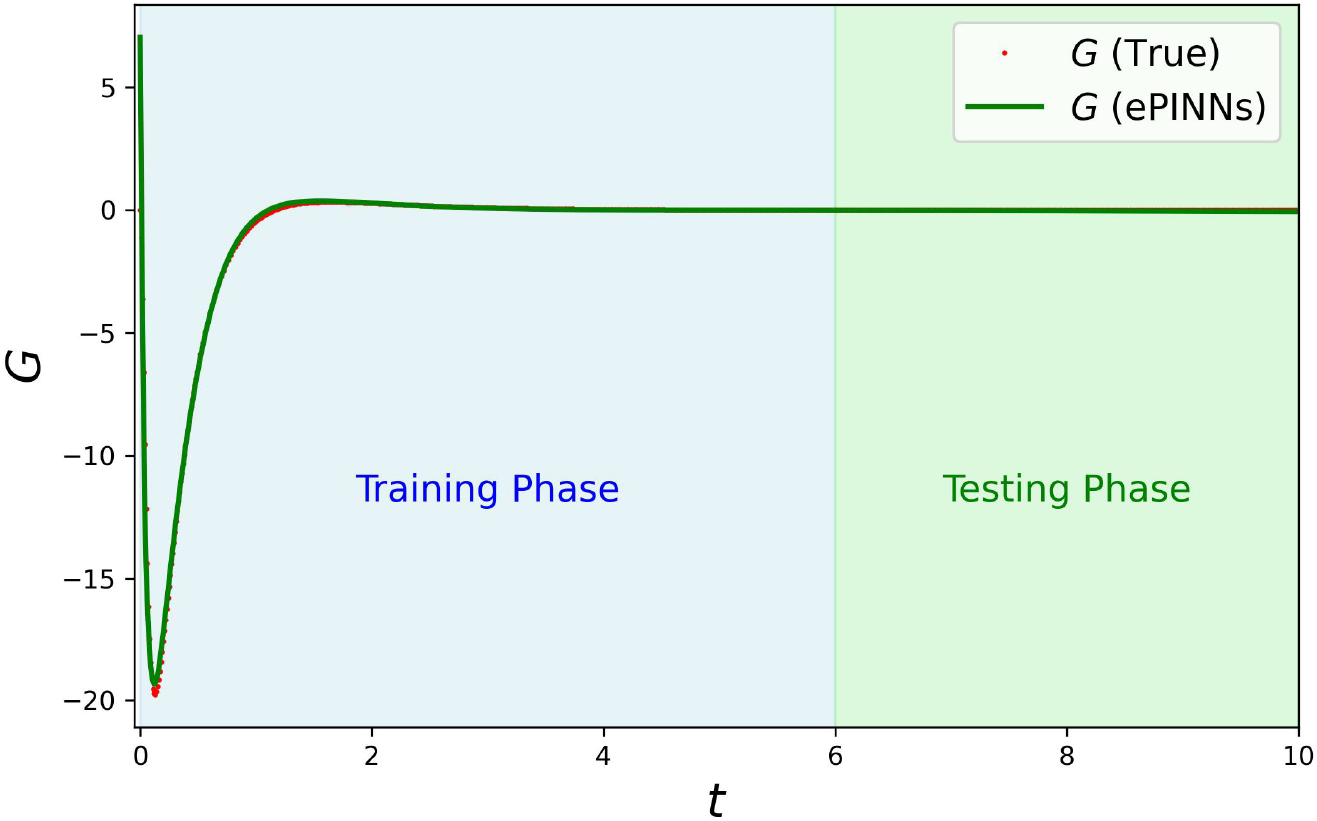
ePINNs prediction of missing dynamics *G*(*x*) with partial data. The solid green line (labeled “ePINNs”) represents the model’s prediction for the unknown dynamics, while red dots (labeled “True”) indicate ground truth values. The training uses noisy observations from *Y, M*, and *M*_*I*_, demon-strating the network’s ability to infer hidden system components.

### 3.3. A result about PUMS identifiability

#### Theorem 1.

A PUMS is identifiable if and only if col *D* ⊆ col *C*

Here, col *M* denotes the column span of a matrix *M*. Notice that asking that col *D* ⊆ col *C* is equivalent to asking that there exists a matrix *A* such that *D* = *CA*. So our example is an identifiable structure in this sense.

**Proof**. Suppose that *D* = *CA*. Take functions as above, so we have that

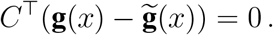

Since *D*^⊤^ = *A*^⊤^*C*^⊤^, it follows that

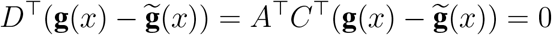

and substituting this into

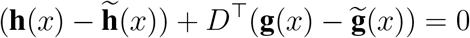

we conclude that 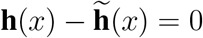.

To prove the converse implication, suppose that col *D* is not included in col *C*. This means that there is a column of *D* which is not in the span of the columns of *C*. Denote the columns of *D* as **d**_*i*_ = (*d*_1*i*_, …, *d*_*ri*_)^⊤^, for *i* = 1, …, *q*. Suppose that **d**_*j*_∉ col *C* for some *j*. Then there exists a vector *ν* ∈ ℝ ^*r*^ in *C*^⊥^ (so that we have *C*^⊤^*ν* = 0) such that *ν*^⊤^**d**_*j*_ ≠ 0. Now pick any **g** and **h** (for example **g** = 0 and **h** = 0). We need to find functions 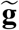 and 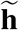 such that (omitting arguments for simplicity): (1) 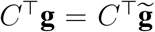 and (2) 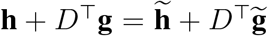, but 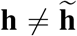. Define 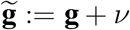 (as a constant function) and 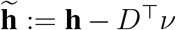. Note that *D*^⊤^*ν*≠ 0 since *ν*^⊤^**d**_*j*_≠ 0, so 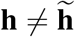. We have that 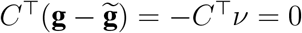, so (1) holds. From

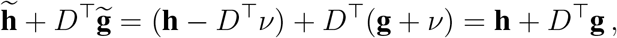

we have that (2) holds as well. This completes the proof.

## 4. DISCUSSION

In this work, we introduced the embedded Physics-Informed Neural Networks (ePINNs) framework to integrate mechanistic modeling with data-driven learning for systems with incomplete knowledge. By combining known system dynamics with neural network-based inference, ePINNs address challenges in modeling biological systems where reaction mechanisms or system parameters are partially unknown.

Applying ePINNs to the study of resource competition in synthetic biology, we demonstrated, through a particular example, their potential ability to infer unmeasured reaction rates from noisy and incomplete observations, predict system behavior beyond observed data using governing ODE constraints, and identify missing dynamics without prior explicit knowledge. Our results showed that physics-informed learning improves estimation accuracy compared to purely data-driven models, particularly for unobserved states. The framework effectively leverages available mechanistic insights while learning unknown interactions, making it adaptable to a broad range of complex biological problems.

In ongoing work, we are evaluating the approach on the more complete resource competition model as well as its performance with other parameter values.

## Acknowledgments

This work was supported in part by grants AFOSR FA9550-22-1-0316 and ONR N00014-21-1-2431, as well as funding from the Institute for Experiential AI at Northeastern University.

## Software

Programs used to generate figures in this report are placed in: *https://github.com/sontaglab/epinn*

## Notes

### Competing Interest Statement

The authors have declared no competing interest.

### Summary of Updates

Typos fixed; 7D model results plotted; some more discussion of identifiability

